# A specific anti-IFITM2 antibody bars the way to SARS-CoV-2 entry into host cells

**DOI:** 10.1101/2022.08.04.502768

**Authors:** Anna Basile, Carla Zannella, Margot De Marco, Gianluigi Franci, Massimiliano Galdiero, Giuseppina Sanna, Aldo Manzin, Massimiliano Chetta, Maria Caterina Turco, Alessandra Rosati, Liberato Marzullo

## Abstract

The early steps of viral infection involve protein complexes and structural lipid rearrangements, which mark the characteristic strategies of each virus in entering permissive host cells. Human IFITM proteins have been described as inhibitors of a broad range of viruses. Despite their homology and functional redundancy, recently it has been surprisingly shown that SARS-CoV-2 is able to specifically hijack the IFITM2 protein. Here has been reported the characterization of a newly generated specific anti-IFITM2 mAb able to impair SARS-CoV-2 Spike protein internalization and, consequently, to reduce the SARS-CoV-2 cytopathic effects and syncytia formation. Importantly, as evidence of the more general involvement of IFITM2 in virus entry, the anti-IFITM2 mAb was able to efficiently reduce HSVs- and RSV-dependent cytopathic effects. Hence, IFITM proteins could be promising targets that can foster the development of biological antiviral molecules, or suggest additional therapeutic strategies for the treatment of viral infections.

**GRAPHICAL ABSTRACT:** 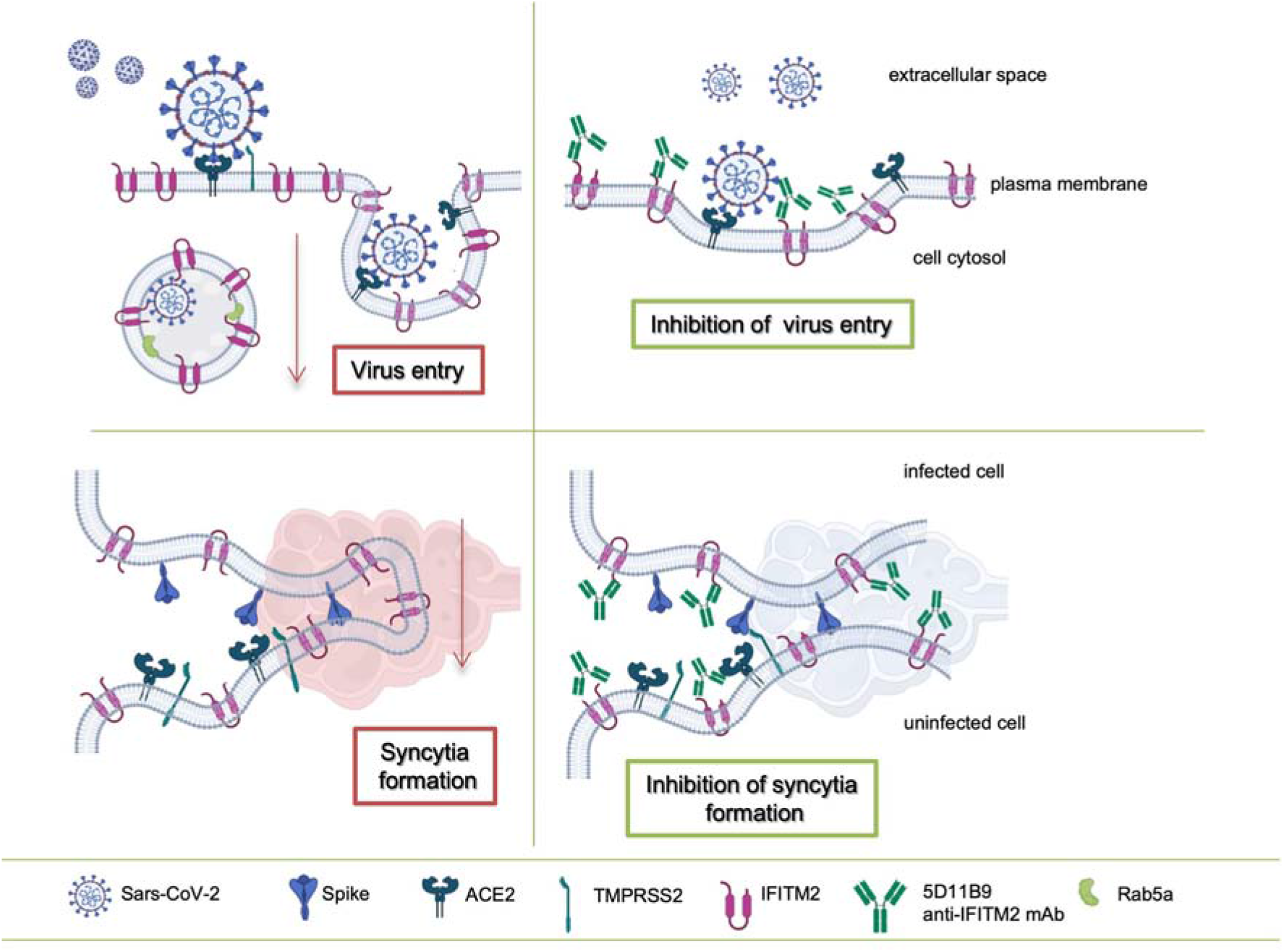

## INTRODUCTION

The severe acute respiratory syndrome coronavirus 2 (SARS-CoV-2) recently turned out to be exceptionally contagious, and the World Health Organization (WHO) classified COVID-19, the SARS-CoV-2-related disease, as a global pandemic on March 11th, 2020. Thereafter, the virus spread to more than 200 countries, with severe public health and economic consequences. From October 2020, the emergence of new strains has been linked to new pathogenic variants in the Spike viral protein, including sense and missense mutations, deletions, and insertions (Korber et al., 2020). The WHO has proposed categorizing variants as Variants of Interest (VOI) and Variants of Concern (VOC), the latter characterized by increased transmissibility and risk of hospitalizations or deaths (Forster et al., 2020; Galloway et al., 2021). The scientific community greatly contributed to pandemic control by paving the way for the development of several safe and effective COVID-19 vaccines (Van Kerkhove, 2021) having a major impact on avoiding mortality and helping countries’ economies return to normal (Ledford, 2022a). Now, we have left behind a two-year pandemic that continues to pose a serious threat to public health in the near future; indeed, emerging virus variants may not respond to current preventive and therapeutic measures that need rapid adjustments and reformulations. While vaccination remains the most important tool in prevention, new medications can fill in where vaccines fail, such as against new variants. Current strategies against SARS-CoV-2 are focused on the development of vaccines or monoclonal antibodies, both targeted at viral determinants to neutralize the infectious agent. Conversely - where applicable - an alternative strategy could exploit the neutralization of determinants on the host cell surface that modulate the viral entry. In this respect, some of the new therapeutics were chosen for their ability to block human proteins rather than viral proteins that SARS-CoV-2 uses to infiltrate cells (Ledford, 2022b) and experts are still trying to figure out how this novel coronavirus attacks the body and decipher mechanisms that point to immune system overreaction with fatal consequences (Mulder, 2022; Baggen et al., 2021). Hijacking IFITM human proteins is one of the possible mechanisms by which SARS-CoV-2 could efficiently infect a wide range of host human cells. IFITM proteins were discovered more than 20 years ago during a screening for proteins induced by Interferon, but only in 2009 it was shown their activity as viral restriction factors able to confer basal and IFN-induced resistance to Influenza A virus and to Flaviviruses (Brass et al., 2009). It was previously reported that IFITM proteins restricted infection mediated by human coronaviruses including SARS-CoV-1 (Huang et al., 2011). In contrast, Bozzo et al. showed that endogenous IFITM proteins were essential for efficient infection and replication of SARS-CoV-2 in various types of human cells. In particular, they showed that IFITM proteins are entry cofactors of SARS-CoV-2 in a way that mimicking peptides and/or commercially available antibodies inhibited SARS-CoV-2 infection of human lung, heart and gut cells (Bozzo et al., 2021). Furthermore, they demonstrated that several SARS-CoV-2 VOCs are dependent on IFITM2 for efficient replication, suggesting that IFITM proteins play a key role in viral transmission and pathogenicity (Nchioua et al., 2022).

The design of a specific monoclonal antibody capable of binding the extracellular N-terminal domain of the IFITM2 protein was detailed in this report, and its activity in hampering SARS-CoV-2 penetration mechanisms was demonstrated. Furthermore, the antibody also inhibited cytotoxic effects induced by herpesvirus or respiratory syncytial viruses, indicating that IFITM2 activity in viral infection affects not only SARS-CoV-2 but also other human pathogenic viruses. The pathway mediated by IFITM2 is therefore a general mechanism operated by different viruses and can represent a versatile target for prevention and therapy.

## RESULTS

### Design and binding characterization of a specific anti-IFITM2 mAb

The high similarity of the amino acid sequences of human IFITM2 and IFITM3 - and to a lesser extent also IFITM1 - posed a first challenging task in the selection of the immunogens able to produce efficient and specific monoclonal antibodies selectively targeting IFITM2 among the other homologous members of the IFITM protein family. Moreover, also the debated topology of the members of these membrane proteins contributed further complexity considered in the preliminary setup of the experiments. Finally, the sequence analyses and the available literature data prompted to work on the hypothesis of type III TM topology for IFITM2 and IFITM3 proteins. Type III TM topology is characterized by the cell surface exposure of both N- and C-termini (NTD and CTD), respectively linked to two antiparallel transmembrane domains (TM1 and TM2) connected to a short cytosolic inner loop (CIL) (Sun et al., 2020; Weston et al., 2014). Due to their intracellular and intramembrane localization, or their very short length, CIL, TMs and CTD (128-132 aa) were not considered in the selection of the immunogen. The sequence alignment of human IFITM2 and IFITM3 NTDs allowed us to identify a short sequence segment encompassing the highest number of unmatching amino acids that was selected as a potential antigen (Fig. 1A). The peptide was chemically synthesized and used for the mice’s standard immunization procedures. The screening of hybridomas allowed the isolation of the clone 5D11B9 that showed the best performance in selectively recognizing IFITM2. Namely, IFITM2 or IFITM3 peptides and IFITM1, IFITM2, or IFITM3 recombinant human proteins were used to set up an indirect ELISA, in order to test 5D11B9 mAb binding performance in comparison with commercially available anti-IFITM2 monoclonal or polyclonal antibodies. Fig 1B shows the results of the analyses confirming the selectivity of 5D11B9 mAb towards both IFITM2 peptide and full length human IFITM2 recombinant protein, while the compared monoclonal anti-IFITM2 antibodies showed cross reactivity against IFITM2 and IFITM3 proteins, as well as the rabbit polyclonal anti-IFITM2 antibody showed a wider cross reactivity against all the human IFITMs tested. Furthermore, no commercial anti-IFITM2 antibody was able to bind the immunogen used to produce 5D11B9 mAb (IFITM2 peptide). To further characterize the 5D11B9 mAb selectivity, the antibodies were tested by Western blot analysis using the recombinant N-terminal domain of IFITM2 and IFITM3 expressed in *Escherichia coli* (*E. coli*) as avidin-tagged peptides. Fig. 1C shows a clear selective reactivity of 5D11B9 mAb against IFITM2 NTD, and no cross reactivity with IFITM3 NTD protein, thus confirming the results obtained by the ELISA assay.

**Figure 1.**
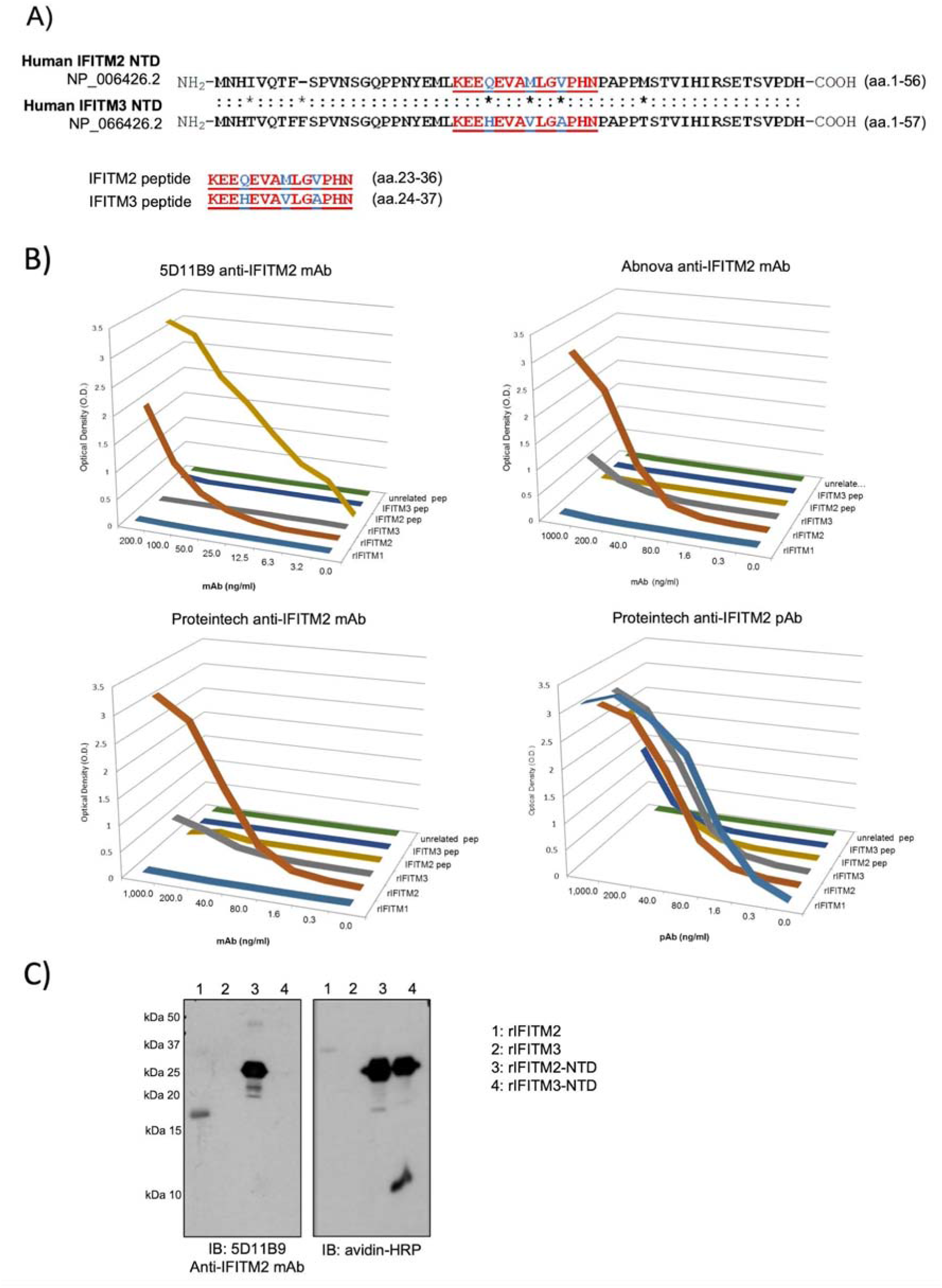
Design and binding characterization of a specific anti-IFITM2 mAb. **A-** Analysis of the N-Terminal (NTD) sequences of IFITM-2 and IFITM-3 and selection of short sequence stretches encompassing the highest number of amino acid substitutions to synthesize oligopeptides for mouse immunization. The sequence alignment and antigen sequences used to immunize the mice are displayed. **B-** IFITM2 peptide, IFITM3 peptide, an unrelated peptide, recombinant human IFITM1, recombinant human IFITM2 and recombinant human IFITM3 were used in the ELISA test. The 5D11B9 anti-IFITM2 mAb binding activity was compared to commercially available anti-IFITM2 antibodies: Abnova anti-IFITM2 mAb, Proteintech anti-IFITM2 mAb and Proteintech anti-IFITM2 pAb. Results were expressed in O.D. means. **C-** Recombinant human IFITM2 and recombinant human IFITM3 proteins, IFITM2 and IFITM3 NTD recombinant peptides were used in the Western blot analyses by using the 5D11B9 anti-IFITM2 mAb. Avidin HRP was used as a positive control for IFITM2 NTD and IFITM3 NTD recombinant peptides.

### The 5D11B9 mAb binds the NTD extracellular domain of plasma membrane-associated IFITM2

Non-permeabilized monkey Vero E6 and human Calu-3 cells were analyzed by flow cytometry in order to check the binding activity of the 5D11B9 anti-IFITM2 mAb and verify the type III TM topology of IFITM2 on the plasma membrane. Vero E6 and Calu-3 cells have been widely used as cellular models for SARS-CoV-2 in *in vitro* studies due to their susceptibility to the virus and the endogenous expression of the proteins ACE2 and TMPRSS2 on the plasma membrane (Lee et al., 2022) (Shang et al., 2020). The cytofluorimetric analysis showed that the 5D11B9 anti-IFITM2 mAb was able to bind the cell surface of both cell lines in a dose-dependent manner (Fig. 2A). On the other hand, confocal microscopy carried out on non-permeabilized Calu-3 cells (Fig 2B) showed that IFITM2 signals are scattered on the whole cell membranes and suggests an uneven distribution of protein clusters. The signal specificity was verified by challenging the binding with competing IFITM2 and IFITM3 peptides. Fig.2C shows that only the IFITM2 peptide is able to compete with 5D11B9 mAb in binding on the Vero E6 and Calu-3 cell surface (p<0.001).

**Figure 2.**
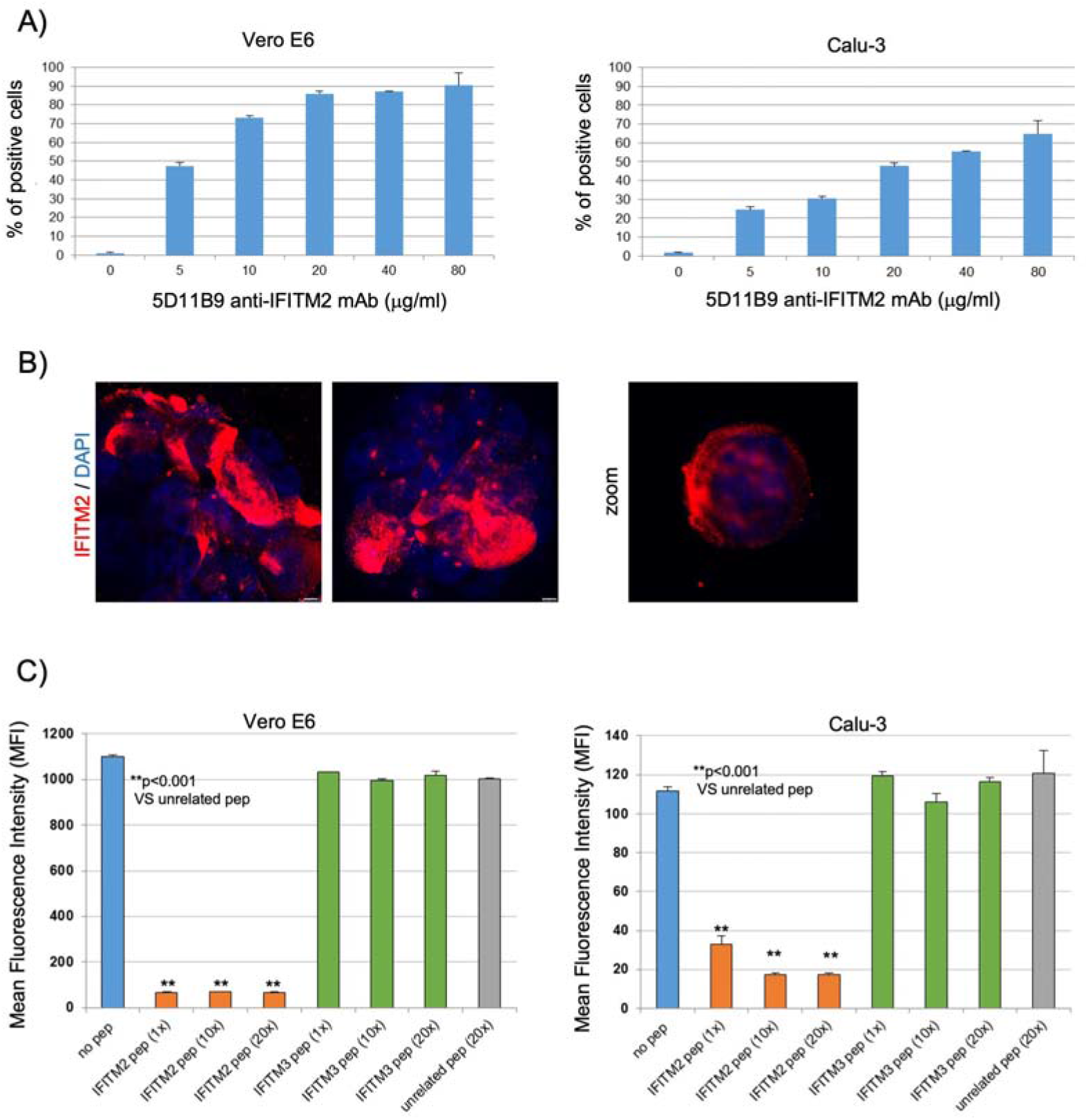
The 5D11B9 mAb binds the NTD extracellular domain of plasma membrane-associated IFITM2. **A-** The monoclonal antibody 5D11B9 was tested for binding to Vero E6 and Calu-3 cell surfaces by flow cytometry. The results showed mAb signal titration from 80 μg/ml to 5 μg/ml as the mean percentage of positive cells (error bars indicate S.D.). **B-** Calu-3 cells were seeded on glass coverslips, processed for immunofluorescence, and analyzed by confocal imaging. Merged images show DAPI staining of DNA (blue) and IFITM2 (red). Scale bars: 5 μm. **C-** In peptide competition assay, the monoclonal antibody 5D11B9 was tested in flow cytometry for binding to the Vero E6 and Calu-3 cell surfaces in the presence of competing peptides (IFITM2 and IFITM3 peptides). Results are represented as Mean Fluorescent Intensity (error bars indicate S.D.). A two-tailed t-test was performed between the indicated groups.

### The 5D11B9 anti-IFITM2 mAb inhibits SARS-CoV-2-Spike protein internalization in host cells

To verify the reported involvement of IFITM2 in the SARS-CoV-2 cell entry process mediated by the binding of the viral Spike (S) protein to the ACE2 receptor, we studied the effects of 5D11B9 mAb treatment on the S-protein internalization. Vero E6 cells were treated with a biotin-tagged recombinant Spike protein, then PE-tagged avidin was used to quantify the membrane bound S-protein. For the assay, human neutralizing and non-neutralizing sera were respectively used as positive and negative controls. A neutralizing serum obtained from SARS-CoV-2 immunized patients, and containing anti-Spike antibodies, efficiently prevented the binding of S-protein onto the cell surface, whereas 5D11B9 mAb considerably increased the binding signal (Fig. 3A), thus indicating the accumulation of S-protein on the cell surface and suggesting the contemporary blockade of the internalization process. Further evidence was obtained by confocal microscopy in Calu-3 cells, as shown in Fig. 3B. Spike signals appear on the cell membrane after 5 minutes of incubation at 37°C, and move towards the cell cytoplasm as soon as S-protein is internalized in about 30 minutes (Fig. 3B, right column: unrelated murine mAb treatment). Conversely, the treatment with the 5D11B9 mAb does not allow S-protein to move into the cytoplasm, as shown by the unaltered signal localization onto the cell surface after 30 min (Fig. 3B, left column). Further evidence of the impairing effects of 5D11B9 anti-IFITM2 mAb treatment on S-protein trafficking was obtained by Proximity-Ligation Assay performed on S-protein, ACE2 (cell surface localization) and Rab5a (endosomal localization) (Bozzo et al., 2021). The assay allowed to confirm the close proximity of S-protein and ACE2 after 5 min of treatment of Calu-3 cells with mouse Fc-tagged S-protein, and contemporary no PLA signal with the endosomal marker Rab5a. After 30 min, the increased PLA signal with Rab5a showed that S-protein was internalized in endosomes, either in control cells or in cells treated with unrelated murine mAbs. At the same time point, the treatment with the 5D11B9 anti-IFITM2 mAb reduced the colocalization of S-protein with Rab5, as demonstrated by the decrease of about 50% of the Rab5a PLA signal, and the equivalent signal persistence of the ACE2 PLA signal related to the membrane localization of S-protein. These pieces of evidence let infer that the 5D11B9 mAb is able to impair the S-protein internalization (Fig. 3C-D), while having little or no effect on ACE2 functions (Supplementary Fig. 1A).

**Figure 3.**
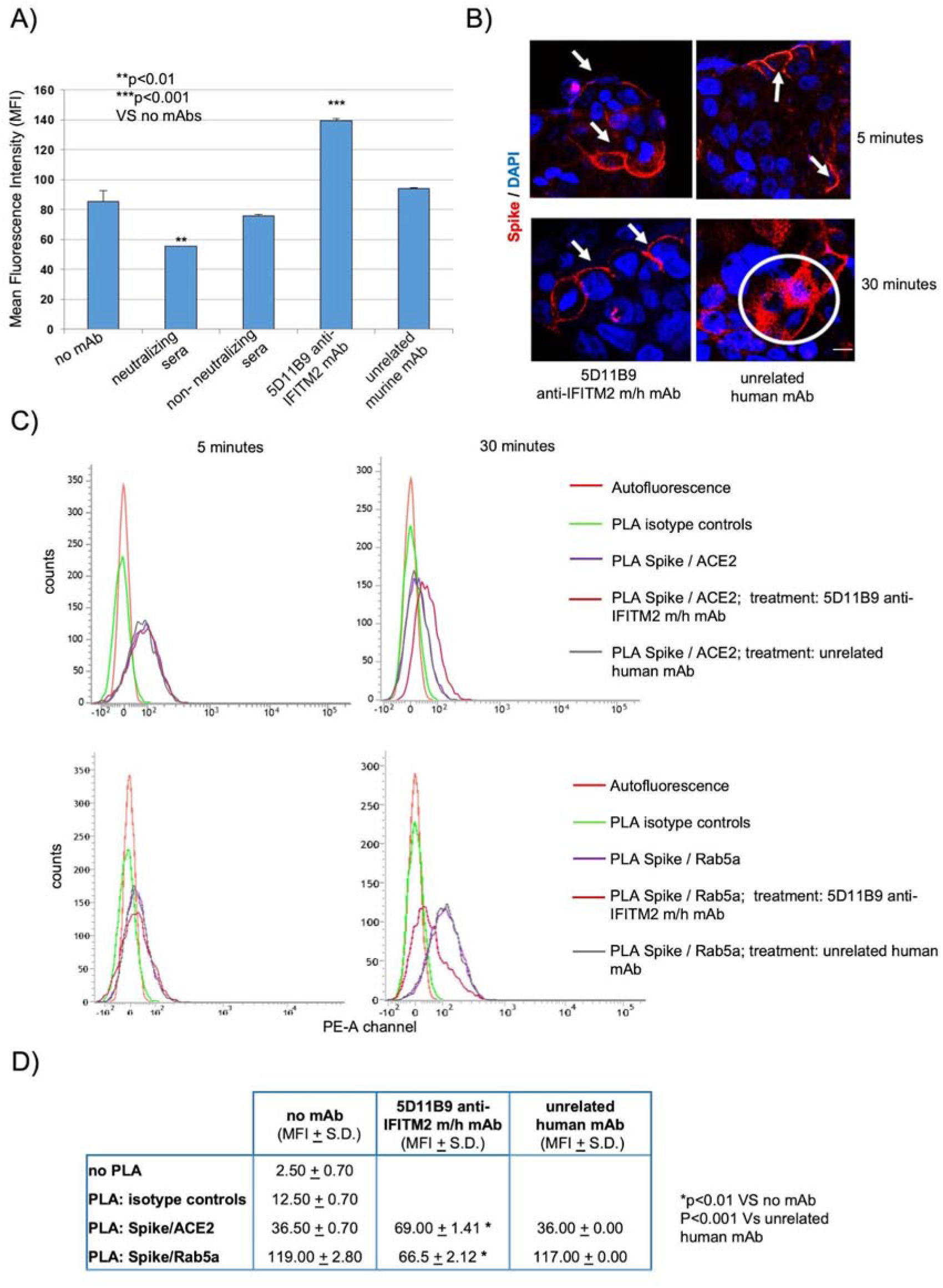
The 5D11B9 anti-IFITM2 mAb inhibits SARS-CoV-2-Spike protein internalization in host cells. **A-** Flow cytometry assays were performed to test the binding of the recombinant biotinylated Spike protein to the Vero E6 cell surface using a PE-conjugated streptavidin. Two human-neutralizing and a non-neutralizing sera (1:30 dilution) were used as positive and negative controls, respectively. The 5D11B9 mAb was added at a concentration of 30 μg/ml. The graph shows the results as Mean Fluorescent Intensity (error bars indicate S.D.). A twotailed t-test was performed between the indicated groups. **B-** Calu-3 cells were treated with a mouse Fc-tagged SARS-CoV-2 S1 spike recombinant protein (5 μg/ml) in the presence of the recombinant 5D11B9 anti-IFITM2 m/h chimera mAb or an unrelated recombinant human mAb (15 μg/ml) for 1 hour at 4°C. Cells were then incubated at 37°C and harvested after 5 and 30 minutes, and the signal from the spike protein was detected by immunofluorescence and confocal microscopy acquisition. The arrows in the figure refer to Spike protein associated with the plasma membrane, while the circle indicates spike protein signal diffusion in the cytoplasm. Merged images show DNA staining with DAPI (blue) and mouse Fc-tagged SARS-CoV-2 S1 spike recombinant protein (red). Scale bar: 5 μm. **C-** Calu-3 cells were treated with a mouse Fc-tagged SARS-CoV-2 S1 spike recombinant protein in the presence of a recombinant 5D11B9 m/h chimera or an unrelated recombinant human mAb for 1 hour at 4°C. Cells were then incubated at 37°C and harvested after 5 and 30 minutes. (UP) PLA signal between Spike and ACE-2 after 5 and 30 minutes at 37°C. (DOWN) PLA signal between Spike and Rab5a after 5 and 30 minutes at 37°C. Graphs depict the signal overlays in the described experimental conditions. **D-** Data obtained from 30 minutes incubation were displayed in a table as Mean Fluorescent Intensity (error bars indicate S.D.). A two-tailed t-test was performed between the indicated groups.

### The 5D11B9 anti-IFITM2 mAb inhibits SARS-CoV-2-induced cytopathic effects and syncytia formation

The data obtained by preliminary *in vitro* analyses suggested further investigations into the efficacy of the anti-IFITM2 mAbs treatment in preventing viral infections of host cells. The 5D11B9 anti-IFITM2 mAb was then used to perform a plaque assay on Vero E6 cells infected with genuine SARS-CoV-2 virus. The efficacy of 5D11B9 mAb treatment was evaluated in comparison with other anti-IFITM2 mAbs commercially available. At 48 hours after co-administration of the virus and a 5D11B9 mAb non-cytotoxic concentration (Supplementary Fig. 1B), virus-induced cytopathic effect was evaluated. The treatment with the 5D11B9 anti-IFITM2 mAb demonstrated its efficacy in reducing the plaques’ number by about 30% (p<0.01), while other mAbs didn’t show any significant effect (Fig. 4A). Furthermore, the treatment appeared to significantly lower the expression of SARS-CoV-2-related genes (namely S and N genes), quantified by RT-PCR in Calu-3 cells in the same experimental settings (Fig. 4B). A yield reduction assay on Vero E6 cells, performed by using supernatants from the experiment shown in Fig. 4B, reported that a 30 μg/ml dose of 5D11B9 mAb, is capable of significantly (p<0.001) reducing by about two orders of magnitude (100x) the viral titer. Severe COVID-19 cases are marked by a major morphological signature represented by the extensive presence of infected multinucleated syncytial pneumocytes associated with lung damage. IFITM proteins - mainly IFITM1 - were also described as inhibitors of Spike-mediated cellular fusion during SARS-CoV-2 infection (Buchrieser et al., 2020). As the 5D11B9 mAb exerted its activity by blocking Spike protein internalization, a GFP split assay on Vero E6 and Calu-3 cells was set up to measure the mAb’s ability to prevent syncytia formation (Fig. 4D). In detail, Spike-expressing donor cells (co-transfected with GFP1-10 plasmid) were mixed at 1:1 ratio with acceptor cells (endogenously expressing ACE2 receptor and co-transfected with GFP11 plasmid), and after 24 hours GFP-positive syncytia were detected and counted. Figure 4E shows a representative image of a GFP-positive syncytia in Vero E6 cells. In agreement with the previous results, the addition of the 5D11B9 anti-IFITM2 mAb, 8 hours after transfections, lowered the number of GFP- positive syncytia. The ratio of GFP-positive cells was quantified by flow cytometry. The results reported in Fig. 4F showed that the 5D11B9 anti-IFITM2 mAb was able to significantly reduce the number of syncytia by roughly 50% in a dosedependent manner, both in Vero E6 and in Calu-3 cells. The specificity of the mAbs for the human protein can also explain the observed higher efficacy in Calu-3 cells.

**Figure 4.**
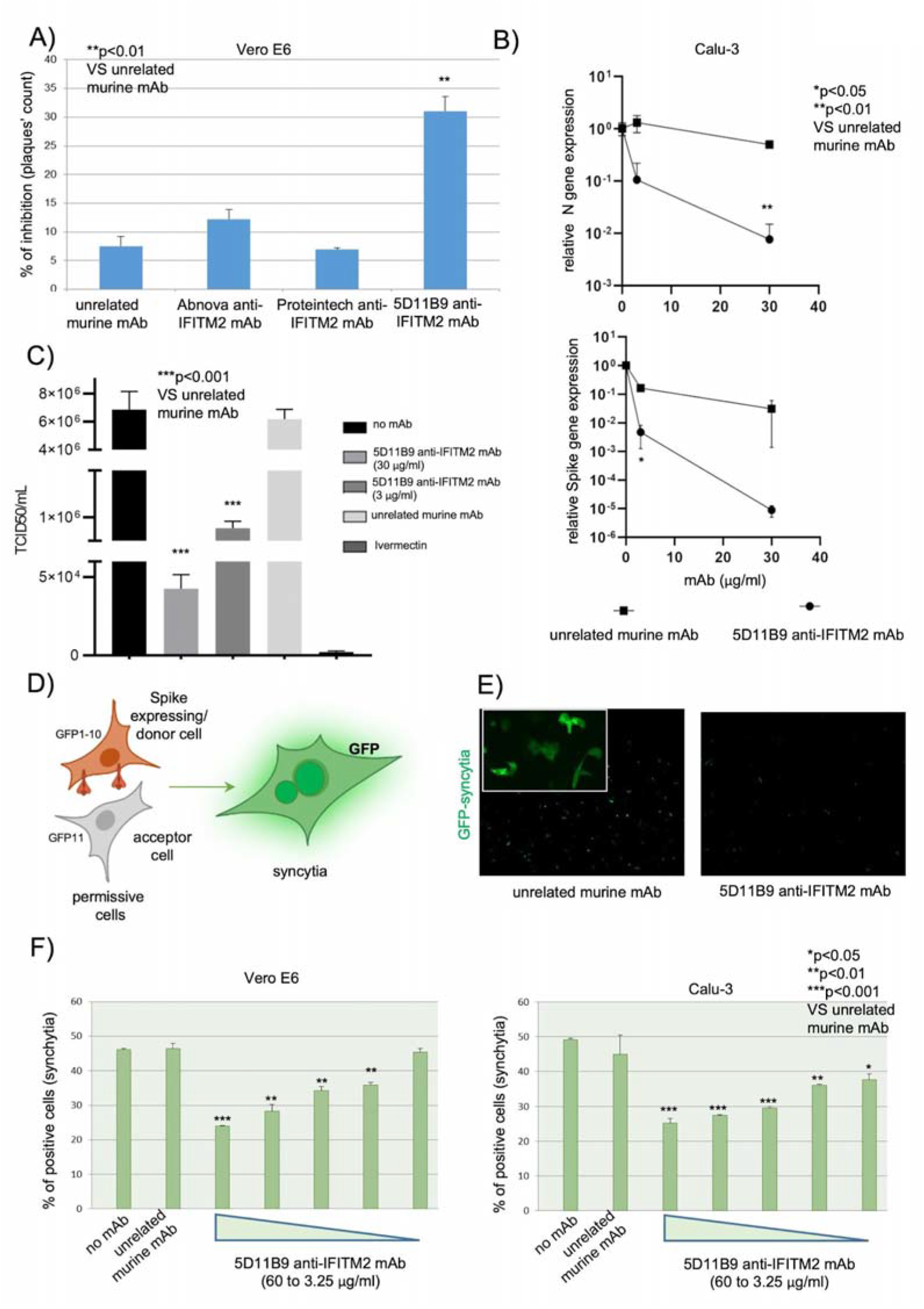
The 5D11B9 anti-IFITM2 mAb inhibits SARS-CoV-2-induced cytopathic effect and syncytia formation. **A-** Vero E6 cells were plated into 12-well cell culture plates (2×10^5^ cells/well) in culture medium. The next day, the monolayer was treated with the 5D11B9 mAb, Abnova anti-IFITM2 mAb, Proteintech anti-IFITM2 mAb or an unrelated murine mAb (30 μg/ml) and simultaneously infected with SARS-CoV-2 (MOI 0.01). After 2 hours, cells were washed with PBS 1X and overlaid with carboxymethylcellulose 0.5% mixed with Dulbecco’s Modified Eagle’s Medium (DMEM). After 48 hrs, the cytopathic effect was first observed under the microscope, and then the plates were stained with 0.5% Crystal Violet / 4% Formaldehyde. The results are shown as a percentage (%) of inhibition of plaque formation (error bars indicate S.D.). A two-tailed t-test was performed between the indicated groups. **B-** Calu-3 cells were plated into 12-well plates (2×10^5^ cells/well) in culture medium. The next day, the monolayer was treated with the 5D11B9 anti-IFITM2 mAb or with an unrelated murine mAb (30 μg/ml or 3 μg/ml) and contemporarily with SARS-CoV-2 (MOI 0.01) at 37°C for 2 hours. After 48 hours, total RNA was obtained and used for Nucleocapsid, Spike and GAPDH gene detection in cultures (supernatants + cells). Graphs depict means (+ SD) calculated as S and N gene relative expression using the 2-^ΔΔCt^ method. GAPDH gene expression was used to normalize RNA level content. **C-** Calu-3 cells were infected with SARS-CoV-2 (100 TCID50). The infected cultures were treated with the 5D11B9 anti-IFITM2 mAb at the indicated doses (30 and 3 μg/ml), and Ivermectin (10 μM) was used as an internal control. Viral yields in the culture supernatant were determined by titration on Vero E6 cells on day 2 post-infection and reported as TCID50/ml. A two-tailed t-test was performed between the indicated groups. **D-** Graphical representation of the GFP-split (syncytia formation) assay developed in Vero E6 and Calu-3 cells. **E-** Representative images of GFP positive syncytia in Vero E6 cells treated with the anti-IFITM2 specific mAb or an unrelated murine mAb. Images are obtained by using 5X magnification. The upper left insert was obtained by using 20X magnification and shows the appearance of multinucleated GFP-positive cells. **F-** Vero E6 (left graph) or Calu-3 (right graph) were transfected with GFP1-10 and Spike or with GFP11 in suspension at 37°C, shaking at 900 rpm for 30 min. After transfection, cells were washed, resuspended in complete medium, and mixed at a 1:1 ratio. After 8 hours, the cells were incubated with different concentrations (60, 30, 15, 7.5, and 3.25 μg/ml) of the 5D11B9 anti-IFITM2 mAb or with an unrelated murine mAb (60 μg/ml). The following day, at 24 hours post-transfection syncytia formation was quantified by flow cytometry and expressed as the mean percentage (%) of positive cells (syncytia). Error bars indicated S.D. and a two-tailed t-test was performed between the indicated groups.

### The 5D11B9 anti-IFITM2 mAb is able to inhibit HSVs- and RSV-induced cytopathic effects

The promising antiviral efficacy resulting from targeting a cellular protein can reverse, where possible, the paradigm of the current research efforts aimed at synthesizing or isolating drugs and/or lead compounds directed against viral components. In fact, like SARS-CoV-2, which exploits ACE2 protein as a docking point on the cell surface, it is often not recommended and potentially dangerous to target cellular proteins that can have pivotal roles in cells or organs’ homeostasis. Although the role of IFITM1-3 deserves a more extensive investigation, their suggested function was mainly restricted in the context of the adaptive immunity and in germ cell specification, thus suggesting low toxicity or side effects risk for a temporary exposure to drugs’ targeting. On the other hand, the specificity of anti-IFITM2 mAbs could potentially reveal their efficacy against other viruses having entry mechanisms into the host cells similar to SARS-CoV-2, or involving IFITM2. In order to investigate the possibility that the 5D11B9 anti-IFITM2 mAb could also impair cell infection by other viruses, we analyzed the effect of the mAb on HSVs- and RSV-replication. Monkey Vero E6 cells and human HSVs-permissive HeLa cells were infected with HSV-1 or HSV-2, and plaques were counted after treatment with the 5D11B9 anti-IFITM2 mAb in comparison with an unrelated murine mAb control treatment. Treatments with 30 μg/ml of the 5D11B9 mAb were able to reduce by approximately 55% the numbers of HSV-1 and HSV-2-induced plaques in Vero E6 cells (Fig. 5A). After the same treatment, a more striking result was observed in human HeLa cells, in which the mAb reduced by about 70% the number of plaques induced by HSV-1 and HSV-2. The observed effect showed a dose-dependency, with a 40 % reduction of plaque counts at 3 μg/ml of the mAb (Fig. 5B), once again suggesting in HeLa cells a greater selectivity of the mAb for the human epitope. Finally, the antiviral efficacy of the 5D11B9 mAb was also tested in the RSV- permissive human HEp-2 cell line (Fig. 5C). A yield reduction assay was performed and the supernatants from human HEp-2 cells, infected with genuine RSV in the presence of the 5D11B9 mAb, or 6-azauridine used as a positive control, were collected and titrated on Vero 76 cells. Notably, a 30 μg/ml dose of 5D11B9 mAb, significantly reduces by about two orders of magnitude (100x) the viral titer.

**Figure 5.**
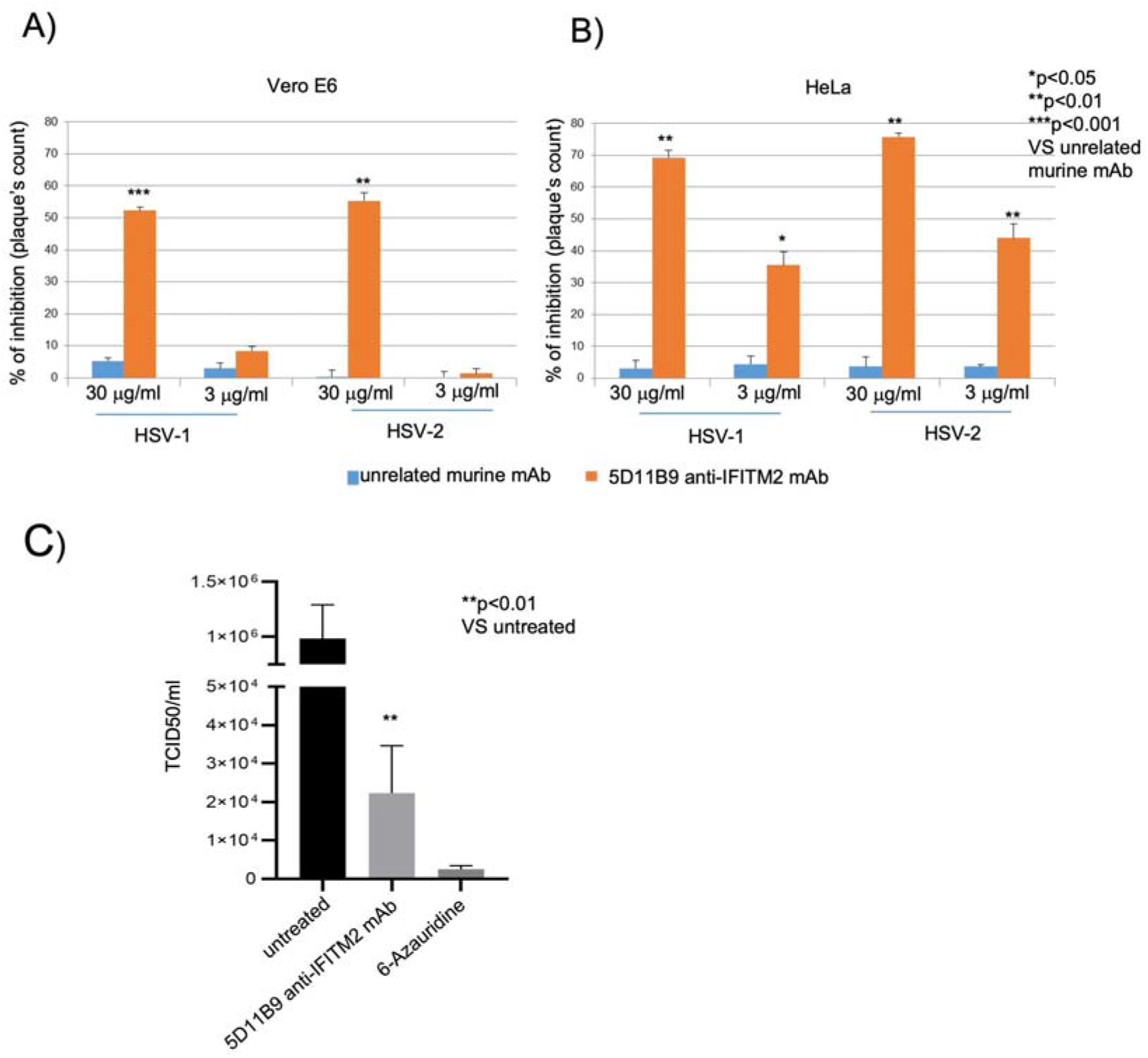
The 5D11B9 anti-IFITM2 mAb is able to inhibit HSVs- and RSV-cytopathic effects. **A,B-** Vero E6 or HeLa cells were plated in 12-well cell culture plates (2×10^5^ cells/well) in culture medium. The next day, the monolayer was treated with the 5D11B9 anti-IFITM2 mAb or an unrelated murine mAb (30 or 3 μg/ml) and simultaneously infected with HSV-1 or HSV-2. Two hours post-infection, cells were washed with PBS 1X and overlaid with carboxymethylcellulose 0.5% mixed with Dulbecco’s Modified Eagle’s Medium (DMEM). After 48h, the cytopathic effect is first observed under the microscope and stained with Crystal Violet 0.5%/Formaldehyde 4%. The results are shown as percentage (%) of inhibition of plaque formation (error bars indicate S.D.) and a two-tailed t-test was performed between the indicated groups. **C-** HEp-2 cells were infected with RSV (100 TCID50). The infected cultures were treated with the 5D11B9 anti-IFITM2 mAb at a concentration of 30 μg/ml. 6-Azauridine (4 μM) was used as an internal control. Viral yields in the culture supernatant were determined by titration in Vero E6 cells on day 4 post-infection and reported as TCID50/ml. Data are expressed as mean (error bars indicate S.D.) and a two-tailed t-test was performed between the indicated groups.

## DISCUSSION

The results illustrated here deepen the knowledge of the IFITM-mediated regulation of cell infection by pathogenic viruses. In particular, the use of a monoclonal antibody capable of selectively recognizing IFITM2 allows it to support the specific role of this member of the IFITM family in mediating cell infection by SARS-CoV-2, HSVs, or RSV.

Although the functional role of IFITM proteins is poorly investigated and still under debate, their dose-dependent modulating effects on the membrane fluidity and curvature have been proposed as triggering the interference/promotion of some virus entry mechanisms. Evidence has also been reported suggesting a role for some IFITM proteins not in allowing, but in restricting the entry of some viruses into the cell (*Diamond and Farzan, 2013*). The availability of reagents that selectively target specific members of the IFITM family is necessary to distinguish the similar or contrasting biological activities of the distinct molecules in different contexts.

To date, immunity to SARS-CoV-2 has been largely obtained in several nations after a massive vaccination campaign, or it has been naturally acquired by people infected by the different SARS-CoV-2 variants that emerged during the pandemic. While vaccines remain the main tool in pandemic control, further complementary therapies are still urgently needed for the prophylaxis and treatment of non-vaccinable people, or subjects whose immune systems do not fully respond to vaccination, or develop breakthrough infections, or simply because of vaccine hesitancy. For these population subsets, it becomes necessary to develop alternative preventive and therapeutic strategies that can be equally effective in containing the contagion and in limiting the damage deriving from the infection, especially in fragile and sensitive subjects (Agrati et al., 2021; Machingaidze & Wiysonge, 2021; Jiménez et al., 2022). The solution proposed in this report overturns, in this case, the classical point of view of the research of the pharmacological targets, from the infectious agent to the cellular determinant/mechanism that allows the latter to finalize the infection process. This reversal of perspective is often difficult or impossible to pursue, since cellular targets and the interference with the processes they regulate entail risks and side effects that are unacceptable and intolerable. Even in the case of SARS-CoV-2 it would be risky to envisage the use of drugs directed against the membrane protein ACE2, used by the virus to anchor itself to the cell and initiate the infection process, since its angiotensin converting activity, through the RAS system, regulates blood pressure with protective functions for the cardiovascular system and other organs (Verano-Braga et al., 2020). On the other hand, we report that IFITM2, although involved in the interaction with the Spike/ACE2 complex, can be bound by a selective mAb without altering the activity of ACE2. This provides an opportunity to block the entry of the virus without unwanted effects. Indeed, to date, no further pieces of evidence showing other fundamental cell functions have been reported for the IFITM proteins, other than their involvement in the innate immune response induced by interferon. These premises reasonably induce us to rely on the absence, or limited, toxicity of anti-IFITM2 treatments with specific monoclonal antibodies, aimed at preventing SARS-CoV-2 infection and subsequent spread.

These pieces of evidence lay the foundations of a hopefully successful, cheaper, and broader spectrum antiviral approach as an alternative or complement to the efforts of a great part of the current industrial investment focused on the production of neutralizing anti-Spike monoclonal antibodies. In this respect, it is worth noting that the efficacy of drugs that target the Spike protein is affected by the changing nature of the protein in increasingly emerging viral variants, which reduces their efficacy and reliability in antiviral therapies. Conversely, as above noted for cellular targets, IFITM2 is not subject to rapid structural variability, and the specific anti-IFITM2 mAb here proposed could be a more durable and widely applicable tool for viral variants, as well as for other viruses exploiting the IFITM2 presence on the plasma membrane and endocytic vesicles. The next steps in developing specific IFITM2 blocking mAbs for potential clinical use will be to start the anti-IFITM2 mAb humanization process and, if necessary, affinity maturation, as well as to validate its activity in viral infection preclinical models. To date, most first line medications do not provide optimal responses, and a growing demand for more valuable virus entry inhibitors is expected in the upcoming COVID-19 post-pandemic era as well as in the context of future emergencies involving new and/or already known viruses.

## Acknowledgements

This study was supported by: University of Salerno Intramural Funds (FARB) to LM; PRIN 2017 “Natural and pharmacological inhibition of the early phase of viral replication (VirSudNet)” N° 2017M8R7N9 to AM and the VALERE project funded by the University of Campania “Luigi Vanvitelli” to MG.

## Author contributions

AR, LM, and MCT designed the studies. AB, CZ, GS, GF, MG, AR, and LM conducted experiments and acquired data. MC, MDM, and AR cured data presentation. AB, MDM, AR, and LM performed statistical analysis. AR, GF, MG, MCT, AM, and LM contributed to manuscript writing and editing.

## STAR⋆Methods

### Key resources table

| REAGENT or RESOURCE | SOURCE | IDENTIFIER |
| --- | --- | --- |
| **Antibodies** |  |  |
| Goat Anti-Mouse IgG Antibody (H&L) [HRP] | Jackson ImmunoResearch | cat# 115-035-003; RRID:AB_10015289 |
| Goat Anti-Rabbit IgG Antibody (H&L) [HRP] pAb | Jackson ImmunoResearch | cat# 111-035-003; RRID:AB_2313567 |
| 5D11B9 murine IgG2b | This paper | N/A |
| recombinant 5D11B9 murine/human IgG4 chimera | This paper | N/A |
| Anti-IFITM2 murine IgG2a | Abnova | cat# H00010581-M14; RRID:AB_1724709 |
| Anti-IFITM2 murine IgG1 | Proteintech | cat# 66137-1-Ig; RRID:AB_2881536 |
| Anti-IFITM2 rabbit IgGs | Proteintech | cat# 12769-1-AP; RRID:AB_2122089 |
| Anti-IFITM2 rabbit IgGs | LsBio | cat# LS-C322156 |
| normal mouse IgG2b-FITC | Santa Cruz Biotechnology | cat#sc-2857; RRID:AB_737258 |
| recombinant unrelated human IgG4 mAb | This paper | N/A |
| recombinant unrelated murine IgG2b mAb | This paper | N/A |
| Anti-mouse IgG2ab-PE | Miltenyi Biotec | cat# 130-097-950; RRID:AB_2661524 |
| Goat Anti-Mouse IgG(H+L) AF555 | SouthernBiotech | cat# 1031-32; RRID:AB_2794317 |
| anti-ACE-2 goat polyclonal | R&D Systems | cat# AF933; RRID:AB_355722 |
| Normal goat IgG | R&D Systems | cat# ab-108-c; RRID:AB_354267 |
| Anti-Rab5a rabbit polyclonal | Santa Cruz Biotechnology | cat#sc-309; RRID:AB_632295 |
| **Bacterial strains** |  |  |
| E. coli | Biotin XCell™ F' cells, Lucigen | Cat# 60704-1 |
| **Chemicals, Peptides, and Recombinant Proteins** |  |  |
| Nitrocellulose membrane | Bio-Rad | Cat#1704159 |
| ECL Western Blotting Substrate | Pierce™, Thermo Fisher Scientific | cat# 32106 |
| recombinant human IFITM1 Protein | Proteintech | cat# Ag2320 |
| recombinant human IFITM2 Protein | Proteintech | cat# Ag17917 |
| recombinant human IFITM3 Protein | Proteintech | cat# Ag17864 |
| Biotinylated Recombinant SARS-CoV-2 Spike protein | R&D Systems | cat# BT10549 |
| SARS-CoV-2 S1 recombinant protein mouse Fc Tag | ProSci | cat# 97-092 |
| Avidin [HRP] | eBioscience™, Thermo Fisher Scientific | cat# 18-4100-94 |
| Streptavidin PE | SouthernBiotech | cat# 7100-09M |
| Gelatin from cold water fish skin | Sigma-Aldrich | cat# G7041 |
| TMB Substrate Solution | Invitrogen, Thermo Fisher Scientific | cat# 00-4201-56 |
| H2SO4 | Sigma-Aldrich | cat# 339741 |
| Nonfat dried milk powder | PanReac AppliChem | cat# A0830 |
| Non-Enzymatic Cell Dissociation Solution | ATCC | cat# 30-2103 |
| FcR Blocking Reagent | Miltenyi Biotec | cat# 130-059-901;<br>RRID:AB_2892112 |
| TransIT X2 Reagent | Mirus Bio LLC | cat# MIR 6005 |
| TRIzol reagent | Sigma-Aldrich | cat# T9424 |
| All-In-One 5X RT MasterMix | Applied Biological Materials | cat# G592 |
| BrightGreen 2X qPCR MasterMix | Applied Biological Materials | cat# G892 |
| Prolong Gold Antifade Mountant | Thermo Fisher Scientific | cat# P36930 |
| Triton X-100 | Sigma-Aldrich | Cat# T9284 |
| Tween 20 | Sigma-Aldrich | Cat# P1379 |
| 0.05 % Trypsin/EDTA (1X) | Gibco | cat# 25300-054 |
| Dulbecco's Phosphate Buffered Saline (PBS) | Corning | cat# 21-031-CV |
| Fetal bovine serum | biosera | cat#FB-1101/100 |
| Penicillin-Streptomycin | Biosera | cat#ZC-A4122/100 |
| Kanamycin | Sigma-Aldrich | Cat#K0245 |
| DAPI (4',6-Diamidino-2-Phenylindole, Dihydrochloride) | Thermo Fisher Scientific | Cat# D1306 |
| Carboxymethylcellulose Low viscosity | Sigma-Aldrich | Cat#C 5678 |
| Carboxymethylcellulose High viscosity | Sigma-Aldrich | Cat# C5013 |
| formaldehyde | Sigma-Aldrich | Cat# CK0272 |
| NaCl | Sigma-Aldrich | Cat# S9625 |
| Tris-HCl | Sigma-Aldrich | Cat# T3253 |
| BSA | Sigma-Aldrich | Cat# A7906 |
| 7-Amino-Actinomycin D | BioLegend | Cat# 420404 |
| **Plasmids** |  |  |
| pAviTag-C N-His SUMO Kan Vector | Lucigen | Cat# 49044-1 |
| pQCXIP-BSR-GFP11 | Addgene | Cat# 68716 |
| pQCXIP-GFP1-10 | Addgene | Cat# 68715 |
| pCMV14-3X-Flag-SARS-CoV-2 S | Addgene | Cat# 68716 |
| **Other** |  |  |
| 96-well microplates | Thermo Fisher Scientific | Cat#442404 |
| SUMO Express Protease | Lucigen | cat#30801-2 |
| NTA-Ni resin | Sigma-Aldrich | cat#P6611 |
| Pierce High-Capacity Endotoxin Removal Spin Column | Pierce | cat#88274 |
| QCL-1000™ Assay | Lonza | cat#50-647U |
| ACE2 Activity Assay Kit Fluorometric | Abcam | cat# ab273297 |
| Duolink In Situ PLA Detection Reagent Far Red | Sigma-Aldrich | cat#DUO92013 |
| Blocking Solution | Sigma-Aldrich | cat#DUO82007 |
| Duolink Antibody Diluent | Sigma-Aldrich | cat#DUO82008 |
| Anti-Mouse MINUS PLA | Sigma-Aldrich | cat#DUO92004 |
| Anti-Rabbit PLUS PLA | Sigma-Aldrich | cat#DUO2002 |
| CCK-8 - Cell Counting Kit-8 | Dojindo | cat#CK04 |

**Table S1:** peptides used in this study.

| Name | Sequence | Description |
| --- | --- | --- |
| IFITM2 peptide | KEEQEVAMLGVPHNC | this study |
| IFITM3 peptide | KEEHEVAVLGAPHNC | this study |
| unrelated peptide | DIDEPRDIPKQPWGPP | this study |

**Table S2.**
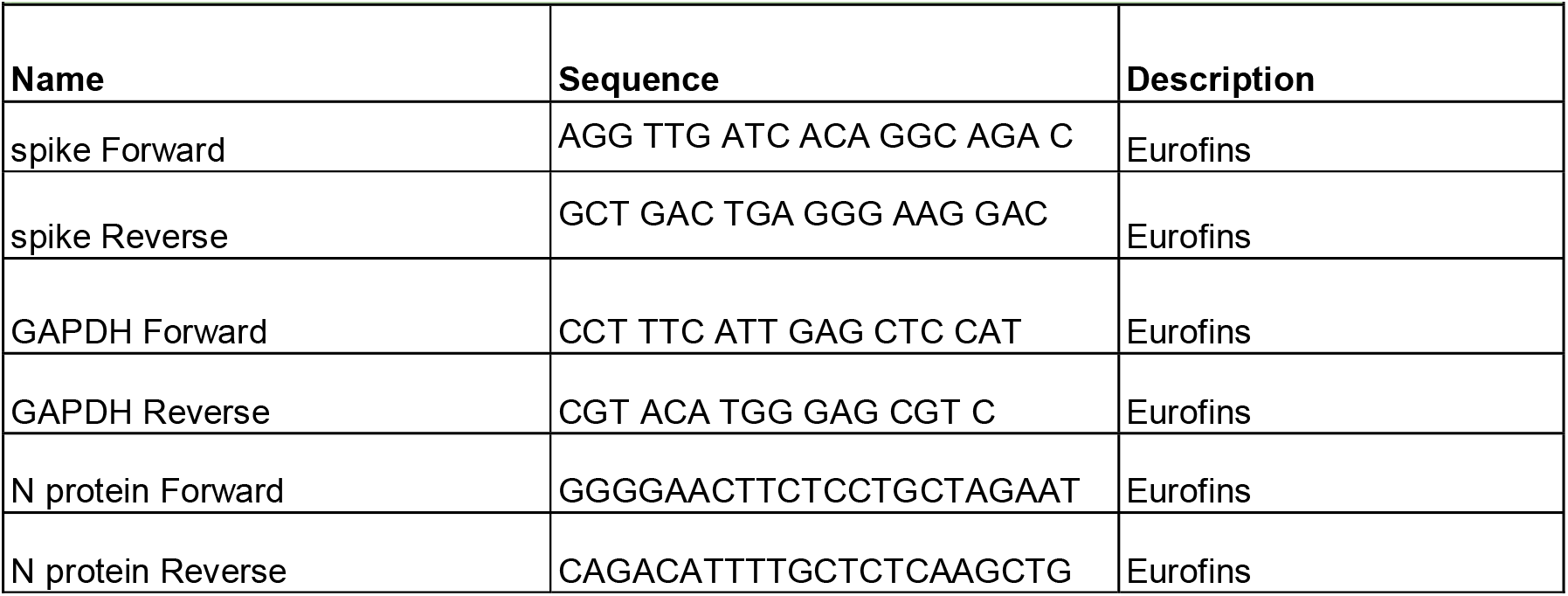
Primers used in the study.

### Resource availability

#### Lead contact

Further information and requests for resources and reagents should be directed to and will be fulfilled by the lead contact, Alessandra Rosati.

#### Materials availability

All unique/stable reagents generated in this study are available from the Lead Contact with a completed Materials Transfer Agreement.

### Experimental models and subject details

#### Virus strains

*SARS-CoV-2 virus -* SARS-CoV-2 (clinical isolate kindly donated by Hospital Lazzaro Spallanzani, Rome, Italy) was propagated in Vero E6 cell monolayers as previously reported in Zannella et al., 2021. *Herpes Simplex (HSV-1 and HSV-2)-* HSV-1 (strain SC16), carrying a lacZ gene driven by the CMV IE-1 promoter to express β-galactosidase (Zannella et al., 2021) and HSV-2 (purchased by ATCC VR-34) were propagated in Vero E6 cell monolayers. *Respiratory Syncytial Virus (RSV)-* Human Respiratory Syncytial Virus (RSV) strain A2 (purchased by ATCC VR-1540) was propagated in Vero-76 cell monolayers (Piras S. et al., 2019).

#### Cell lines

Cell lines were purchased from the American Type Culture Collection (ATCC). The absence of mycoplasma contamination was checked periodically by the Hoechst staining method. Cell lines supporting the multiplication of RNA and DNA viruses were the following: monkey kidney Vero-76 (ATCC CRL 1587 *Cercopithecus aethiops*) or Vero E6 (ATCC CRL-1586 *Cercopithecus aethiops*), human laryngeal carcinoma HEp-2 (ATCC CCL-23), human cervical adenocarcinoma HeLa (ATCC CRM-CCL-2) and human lung adenocarcinoma Calu-3 (ATCC HTB-55). HEp-2 cells were maintained in Minimum Essential Medium with Earle’s salts (MEM-E), L-glutamine, 1 mM sodium pyruvate, and 25 mg/L kanamycin, supplemented with 10% fetal bovine serum (FBS). Vero-76 cells were grown in Dulbecco’s Modified Eagle Medium (D-MEM) with L-glutamine and 25 mg/L kanamycin or PEN/STREP supplemented with 10% FBS. Vero E6 and HeLa cells were grown in Dulbecco’s Modified Eagle Medium (D-MEM) with L-glutamine and PEN/STREP supplemented with 10% FBS. Calu-3 cells were maintained in Eagle’s Minimum Essential Medium salts (EMEM), L-glutamine, 1 mM sodium pyruvate, supplemented with 10% or PEN/STREP supplemented with 10% FBS. Cell cultures were grown in a 37°C incubator with a humidified atmosphere of 5% CO2 in the air.

### Methods details

#### Antibody production

The murine monoclonal 5D11B9 anti-IFITM2 antibody and the recombinant 5D11B9 murine human IgG4 chimera were produced by Genscript. The unrelated murine mAb was provided both in murine IgG2b or human IgG4 format by Evitria.

#### Cloning and expression of recombinant IFITM2 and IFITM3 NTDs

The NTDs (N-Terminal Domains) of IFITM2 (aa 1-56) and IFITM3 (aa 1-57) were cloned in pC-AviTag SUMO Vector™ (Lucigen, WI, USA) and expressed in *E. coli* as fusion protein with biotinylated tag. The expression and production of the proteins were then induced and optimized according to the manufacturer’s instructions. As expected, the recombinant proteins carried a fused C-terminal biotinylated (Bt) tag. The subsequent cleavage with SUMO Express Protease allowed the release of the Bt-tagged NTDs that were subsequently purified on NTA-Ni resin to remove the contaminants (His-SUMO tag and His-tagged protease).The Pierce High-Capacity Endotoxin Removal Spin Column was used to obtain endotoxin-free preparations. Endotoxin concentration was measured by QCL-1000™ Assay following the manufacturer’s instructions.

#### Confocal microscopy

Calu-3 cells were seeded on glass coverslips. After 24 hours, cells were washed in PBS 1X, fixed with 3.7 % formaldehyde in PBS 1X for 20 minutes at room temperature (RT), washed in 0.1 M glycine and blocked in PBS 1X containing 3% BSA for 30 minutes at RT. Cells were then immunostained with the 5D11B9 murine IgG_2b_ mAb (3 μg/ml) overnight at 4°C. The primary antibody was detected with AF55-conjugated secondary anti-mouse IgGs (1:200) for 1 hour at RT. Calu-3 cells were also incubated for 2 hours at 4°C. After that, a recombinant mouse-Fc tagged SARS-CoV-2 S1 protein was added to the culture media at a concentration of 5 μg/ml in combination with a recombinant 5D11B9 m/h IgG_4_ chimera (15 μg/ml) or with an unrelated human IgG_4_ (15 μg/ml). After one additional hour at 4°C, the cells were incubated at 37°C and harvested after 5 and 30 minutes. At the end of treatments, cells were washed once with PBS 1X, fixed with 3.7 % formaldehyde in PBS for 20 minutes at RT, washed in 0.1 M glycine and then permeabilized with 0.005 % saponin in PBS 1X/BSA 3 % for 30 minutes at RT. Then, in order to detect the spike recombinant fused protein, cells were incubated with an AF55-conjugated secondary anti-mouse IgGs (1:200) for 1 hour at RT. Slides were then mounted with Prolong Gold Antifade Mountant as well as with DAPI (4’,6-Diamidino-2-Phenylindole, Dihydrochloride) to visualize nuclei and analyzed using a confocal laser scanning microscope (Leica SP5, Leica Microsystems, Wetzlar, Germany). Images were acquired in sequential scan mode by using the same acquisition parameters (laser intensities, gain photomultipliers, pinhole aperture, and 63X objective) when comparing experimental and control material. For figure preparation, brightness and contrast of images were adjusted by taking care to leave a light cellular fluorescence background for visual appreciation of the lowest fluorescence intensity features and to help comparison among the different experimental groups.

#### ELISA assay for anti-IFITMs antibodies

96-well microplates were coated with 50 μL of solutions containing the following peptides or proteins: IFITM2 peptide, IFITM3 peptide, unrelated peptide, recombinant human IFITM1, recombinant human IFITM2, recombinant human IFITM3 (1 μg/ml in PBS 1X) and incubated overnight at 4 °C. The day after, wells were washed with PBS 1X containing 0.1% Tween (washing buffer) and the blocking of nonspecific sites was performed for 1 h at room temperature in PBS 1X containing 0.5% fish gelatin. Hence, plates were washed five times with the washing buffer and loaded with different concentrations of anti-IFITMs monoclonal and polyclonal antibodies. Plates were then extensively washed and incubated for 30 minutes at room temperature with HRP-conjugated anti-mouse IgGs 1:20,000 or anti-rabbit 1:2,000. Subsequently, the TMB solution 1X was added to the wells for the analyte detection. The chromogenic reaction was blocked by acidification with 0.5 m H_2_SO_4_, and the optical density (O.D.) was measured at 450 nm.

#### FACS binding assay

Vero E6 or Calu-3 cells were washed with PBS 1X and treated with Non-Enzymatic Cell Dissociation Solution. Cells were harvested by centrifugation and incubated with 80 μl of PBS 1X containing 10% of decomplemented FBS and 0.1% NaN_3_ (binding buffer) and 20 μl of FcR Blocking Reagent for 15 min on ice, following the manufacturer’s instructions. Then, cells (1×10^6^/ml for cell lines) were resuspended in binding buffer and incubated for 15 min on ice. Thereafter, cells were incubated with different concentrations of monoclonal 5D11B9 murine mAb in binding buffer (100 μl) for 30 minutes on ice. An unrelated murine mAb was used as negative controls. After incubation, cells were washed three times with PBS 1X containing 2% of decomplemented FBS and 0.1% NaN_3_ (washing buffer), centrifuged for 10 minutes at 300 g, the bound monoclonal antibody was revealed by the addition of 100 μl a 1:50 dilution of PE-conjugated anti-mouse IgG and incubated in binding buffer for 30 min on ice. After incubation, cells were washed three times with a washing buffer, centrifuged for 10 minutes at 300 g, resuspended in 300 μl of binding buffer and analyzed by flow cytometry. 7-Amino-Actinomycin D (7-AAD) was used for the exclusion of nonviable cells in flow cytometric assays.

#### GFP-Split fusion assay

For the GFP-split fusion assay, 1×10^6^ cells were separately transfected with 250 ng of DNA in suspension at 37°C shaking at 900 rpm for 30 min using TransitX2 reagent. Donor cells were transfected with 125 ng of pQCXIP-GFP1-10 and 125 ng pCMV14-3X-Flag-SARS-CoV-2 S. Acceptor cells were transfected with 125 ng of pQCXIP-BSR-GFP11 and 125 ng of an empty vector. After transfection, cells were washed and resuspended in complete medium, mixed at a 1:1 ratio in different combinations, and plated at 2×10^5^ cells per well in a 24-well plate. After 8 hours, cells were incubated with different concentrations of the 5D11B9 mAb (60, 30, 15, 7.5 and 3.25 μg/ml) or with an unrelated murine mAb (60 μg/ml). The next day, 20 hours post-transfection, cells were trypsinized and analyzed by flow cytometry. 7-Amino-Actinomycin D (7-AAD) was used for the exclusion of nonviable cells in flow cytometric assays.

#### Neutralization assay by FACS

Biotinylated recombinant SARS-CoV-2 Spike protein (10 μg/ml) was pre-incubated 30 minutes at room temperature with patient samples of human serum containing–SARS-CoV-2 neutralizing antibodies (1:30) in binding buffer (50 μl). Vero E6 cells were also pre-incubated with monoclonal 5D11B9 murine IgG_2b_ (30 μg/ml) in binding buffer (50 μl) 30 minutes on ice or challenged with human neutralizing or non-neutralizing sera (1:30 dilution). The mix was added to 50 μl of cell suspension (1×10^5^ cells) and, after 1 hour of incubation at 37°C, cells were washed three times with a washing buffer and centrifuged for 10 minutes at 300 g. The bound biotinylated recombinant SARS-CoV-2 Spike protein was revealed by the addition of 100 μl of PE-Streptavidin (1:200) and incubated in binding buffer for 30 min on ice. After incubation, cells were washed three times with washing buffer, centrifuged for 10 minutes at 300 g, resuspended in 300 μl of binding buffer and analyzed by flow cytometry. 7-Amino-Actinomycin D (7-AAD) was used for the exclusion of nonviable cells in flow cytometric assays.

#### Peptide competition assay by FACS

Monoclonal 5D11B9 murine IgG_2b_ (20 μg/ml), were pre-incubated 30 minutes at room temperature with the IFITM2 peptide or IFITM3 peptide or unrelated peptide at different concentrations (1X, 10X and 20X) in binding buffer (50 μl). The mix was added to 50 μl of cell suspension (1×10^6^ cells) and, after 30 min of incubation on ice, cells were washed with a washing buffer and centrifuged for 10 minutes at 300 g. The bound monoclonal antibody was revealed by the addition of a PE-conjugated secondary anti-mIgG_2ab_ (1:50) and incubated in binding buffer for 30 min on ice. After incubation, cells were washed with washing buffer, centrifuged for 10 minutes at 300 g, resuspended in 300 μl of binding buffer and analyzed by flow cytometry. 7-Amino-Actinomycin D (7-AAD) was used for the exclusion of nonviable cells in flow cytometric assays.

#### Plaque assay

Vero E6 cells were plated in 12 multi wells (2×10^5^ cells/well). After 24 hours, cells at 80 % confluence were treated with each Ab and simultaneously infected with the SARS-CoV-2 (at a viral titer of 5×10^6^ PFU/mL) or other types of viruses, with DNA genome i.e. HSV-1 (Herpes Simplex Virus-1 and HSV-2 (Herpes Simplex Virus-2). After the adsorption time of 2h, time necessary for the virus to take root and enter the host cells, plates were washed with 1X PBS, and a mixture of 3 % carboxymethylcellulose/10 % FBS 1:3 culture medium was administered. After 48h, the cytopathic effect was first observed under the microscope and then plates were stained with Crystal Violet 0.5 %/Formaldehyde 4 %. Plaques were counted under the microscope and viral inhibition was calculated against the untreated virus control according to the formula: 100 - [(treated well plaques / plaques in infected control) x 100].

#### RT-PCR

Total RNA was isolated using TRIzol reagent and quantified by measuring the absorbance at 260/280 nm (NanoDrop 2000, Thermo Fisher Scientific, Waltham, MA, USA). Approximately 1 μg of total RNA was reverse-transcribed to cDNA by 5X All-In-One RT MasterMix (Applied Biological Materials, Richmond, Canada). A quantitative polymerase chain reaction was run in triplicate using a CFX Thermal Cycler (Bio-Rad, Hercules, CA, USA). 2 μl of cDNA was amplified in 20 μl reactions using BrightGreen 2X qPCR MasterMix-No Dye (Applied Biological Materials, Richmond, Canada) and 0.1 μM of primer. Relative target Ct (the threshold cycle) values of the S and N genes were normalized to Glyceraldehyde 3-phosphate dehydrogenase (GAPDH), as a housekeeping gene. Relative mRNA levels were expressed using the 2-^ΔΔCt^ method.

#### Proximity Ligation Assay

Calu-3 cells were trypsinized, collected into eppendorf tubes, and centrifuged at 500 *g* for 5 min. The cells in complete medium were then incubated with the recombinant mouse-Fc tagged SARS-CoV-2 S1 protein (5 μg/ml) in combination with the recombinant 5D11B9 murine/human IgG_4_ chimera (15 μg/ml) or with an unrelated human IgG_4_ (15 μg/ml). After one hour at 4°C cells were incubated at 37°C and harvested after 5 and 30 minutes. For analysis of protein–protein interactions using flow cytometry, we used the Duolink In situ PLA Detection Reagent FarRed. Briefly, cells were centrifuged at 500× *g* for 5 min, washed with PBS 1X, and fixed with 3.7 % formaldehyde/PBS 1X for 10 min. After two washes with PBS 1X, the cells were permeabilized with 0.5 % Triton/PBS 1X for 7 min. Subsequently, the cells were blocked in Blocking Solution at 37 °C for one hour. Next, the cells were aliquoted in a 96-well V-bottom plate (100,000 cells per well) and then incubated at 37°C for one hour with primary antibodies, polyclonal Goat antibody ACE-2 (5 μg/ml) or polyclonal Rabbit antibody Rab5a (5 μg/ml), that were diluted in Duolink Antibody Diluent. Following washing in wash buffer A (10 mM Tris, 150 mM NaCl, and 0.05% Tween 20), the cells were incubated with secondary antibodies conjugated with PLUS and MINUS probes (Sigma-Aldrich) for one hour at 37 °C. Next, the cells were again washed twice in wash buffer A and then incubated with the ligase (diluted 1:40 in ligation buffer) for 30 min at 37 °C. Following the next round of washing in wash buffer A, the cells were incubated with the polymerase (diluted 1:80 in amplification buffer) for 16 hours at 37 °C. Finally, the cells were washed in wash buffer B (200 mM Tris and 100 mM NaCl), re-suspended with 300 μL of 2 % ΔFBS in PBS 1X and analyzed by flow cytometry.

#### Western blot analysis

100ng of recombinant human IFITM2 protein or recombinant human IFITM3 protein or IFITM2 NTD peptide or IFITM3 NTD peptide were run on 15% SDS–PAGE gels and electrophoretically transferred to a nitrocellulose membrane. Nitrocellulose blots were blocked with 10% non-fat dry milk in TBST buffer (20 mM Tris–HCl pH 7.4, 500 mM NaCl, and 0.1% Tween 20) and incubated with the primary antibody in TBST containing 5% non-fat dry milk, overnight at 4°C. Immunoreactivity was detected by sequential incubation with HRP-conjugated secondary antibody, (HRP-anti-mouse 1:5,000) or HRP-conjugated Avidin (1:1,000) and ECL reagents.

### Yield reduction assays

Hep-2 cells or Calu-3 cells (5×10^5^/ml) were infected with RSV or SARS-CoV-2 (100 TCID50) respectively in maintenance medium and contemporary mAbs and compounds were tested. After 96 hrs at 37° C and 5% CO_2_, each sample was harvested and stored at −80°C. Samples were then diluted with serial passages, starting from 10^-1^ up to 10^-10^. The titer of the virus-containing supernatant dilutions series was determined by the TCID50 end-point in Vero 76 cells.

### Software and statistical analysis

Graphs were realized by using Excel (Suite Office 365) and GraphPad Prism 8.0. Results are expressed as means ± SD. Data were analyzed by Student’s t-test and *p* values <0.05 were considered statistically significant.

## Supplementary Material

**Supplementary Figure 1.**
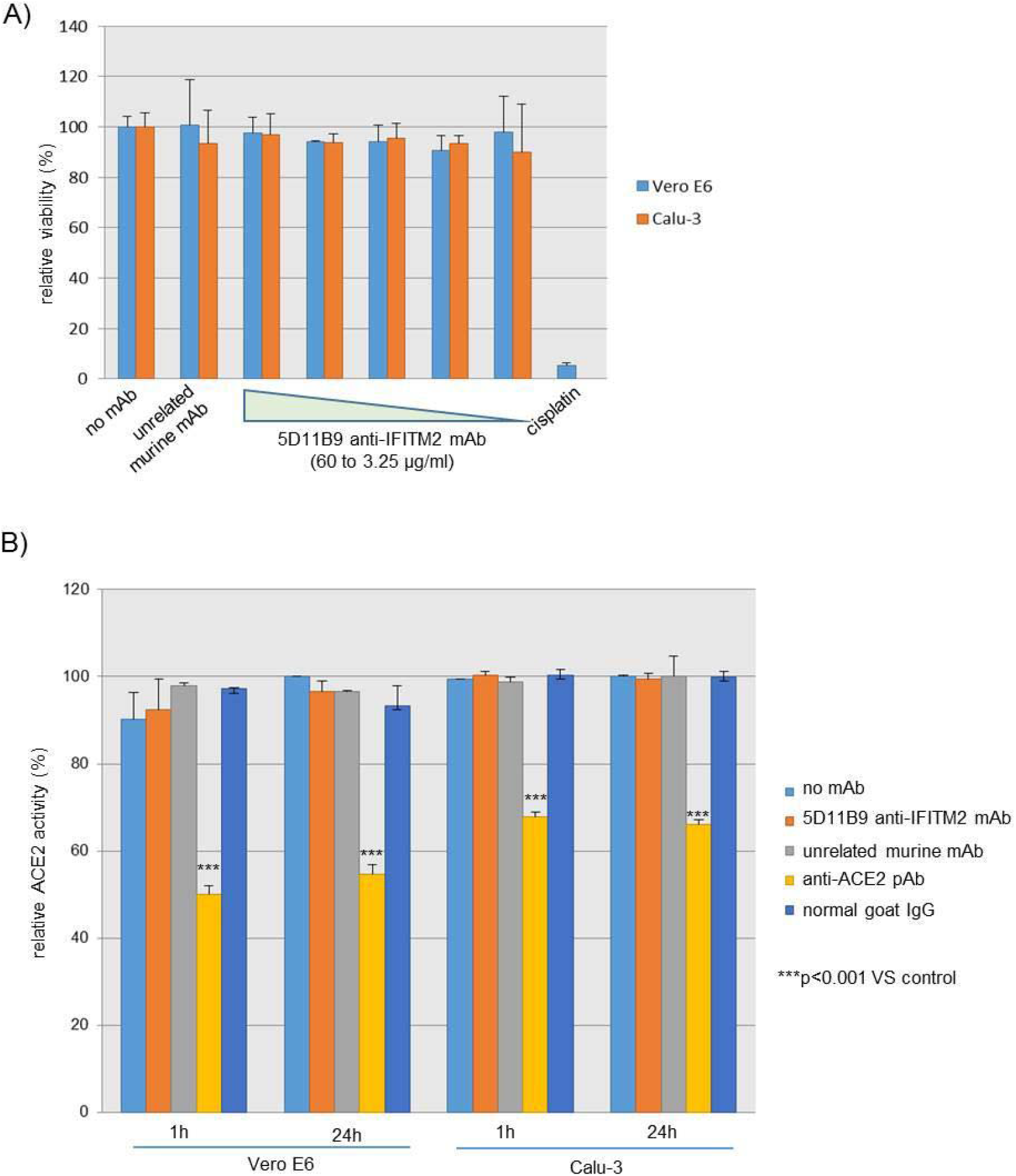
A- Vero E6 and Calu-3 cells were seeded in a 96-well plate at 1×10^5^ cells/well and after an overnight incubation at 37°C cells were treated with the 5D11B9 anti-IFITM2 mAb or with a unrelated murine mAb and (30 μg/ml) for 1 and 24 hours at 37 °C. The anti-ACE2 pAb (10 μg/ml) was used as positive control and normal goat IgG used as a negative control. Cells were then washed with PBS 1X before performing the ACE-2 activity assay. Cells were incubated for 30 min at 37 °C and transferred to black 96-well plate for fluorescence reading (320/420 nm). Results are shown as relative ACE-2 activity (%). Error bars indicate S.D. B- Vero E6 and Calu-3 cells were cultured in 96-well plates and treated with different concentrations of monoclonal antibody 5D11B9 (80, 40, 20, 10 and 5 μg/ml) or with an unrelated murine IgG2b (60 μg/ml). After 48 hours of incubation, cells were analyzed by CCK-8 cell viability assay. Data was obtained from triplicate samples. The results are shown as the percentage (%) of viable cells with respect to control.

## Supplemental Methods

### Ace-2 enzymatic activity assay

The catalytic activities of endogenous ACE-2 were detected by the ACE-2 Activity Assay Kit Fluorometric. Vero E6 and Calu-3 cells were seeded in a 96-well plate at 5×10^4^ cells/well and cultured overnight. The cells were then incubated with 30 μg/ml the 5D11B9 mAb or with 10 μg/ml of the anti-ACE2 pAb or with isotype controls (murine IgG_2b_ and normal goat IgGs) for 1 and 24 hours at 37 °C. Cells were then washed with PBS 1X before the ACE-2 activity assay according to the manufacturer’s instructions. Cells were incubated for 30 min at 37 °C and then transferred to black 96-well plate for fluorescence reading (320/420 nm).

### Cell viability assay

Vero E6 (1×10^4^/well) and Calu-3 (2×10^4^/wel) cells were seeded into the 96-well plates and cultured overnight at 37°C. The cells were then incubated with different concentrations of monoclonal antibody 5D11B9 (80, 40, 20, 10 and 5 μg/ml) or with an unrelated murine IgG_2b_ (60 μg/ml). Cisplatin (30 μM) was used as a known cytotoxic agent. After 48 hours, the CCK-8 solution was added to each well of cells following the manufacturer’s instructions. After 1 hour of incubation, the absorbance of each well was measured using a microplate reader (450 nm), after which the results were statistically analyzed.

